# Brain-Controlled Electrical Stimulation Restores Continuous Finger Function

**DOI:** 10.1101/2022.06.15.496349

**Authors:** Samuel R. Nason-Tomaszewski, Matthew J. Mender, Eric Kennedy, Joris M. Lambrecht, Kevin L. Kilgore, Srinivas Chiravuri, Nishant Ganesh Kumar, Theodore A. Kung, Matthew S. Willsey, Cynthia A. Chestek, Parag G. Patil

## Abstract

Brain-machine interfaces have shown promise in extracting upper extremity movement intention from the thoughts of nonhuman primates and people with tetraplegia. Attempts to restore a user’s own hand and arm function have employed functional electrical stimulation (FES), but most work has restored discrete grasps. Little is known about how well FES can control continuous finger movements. Here, we use a low-power brain-controlled functional electrical stimulation (BCFES) system to restore continuous volitional control of finger positions to a monkey with a temporarily paralyzed hand. In a one-dimensional, continuous, finger-related target acquisition task, the monkey improved his success rate to 83% (1.5s median acquisition time) when using the BCFES system during temporary paralysis from 8.8% (9.5s median acquisition, equivalent to chance) when attempting to use his temporarily paralyzed hand. With two monkeys under general anesthesia, we found FES alone could control the monkeys’ fingers to rapidly reach targets in a median 1.1s but caused oscillation about the target. Finally, when attempting to perform a virtual two-finger continuous target acquisition task in brain-control mode following temporary hand paralysis, we found performance could be completely recovered by executing recalibrated feedback-intention training one time following temporary paralysis. These results suggest that BCFES can restore continuous finger function during temporary paralysis using existing low-power technologies and brain-control may not be the limiting performance factor in a BCFES neuroprosthesis.

## INTRODUCTION

Neural prostheses have the potential to restore function and independence to people with neurological disorders and injuries. By interfacing with the nervous system directly, they can extract a user’s intention information from the native controlling circuits and use those signals to control prostheses, computers, or reanimate paralyzed limbs. Particularly in tetraplegia, it has been found that restoration of hand and arm function is of greatest importance^1^. Furthermore, for the purposes of upper extremity restoration, people with paralysis would prefer use of their natural arm and hand over external prostheses^2^.

Functional electrical stimulation (FES) has shown promise in restoring function to paralyzed arms and hands for reaching and grasping^3–6^. By delivering electrical current to partially or completely non-functional musculature, commercial devices like the Freehand System^7,8^ have already been used with people with upper extremity paralysis to restore movement of native arms and hands. More recently, systems like the Networked Neuroprosthesis (NNP) have shown promise in primarily research-focused studies, having been implanted in five people with spinal cord injury to date^9,10^. In cases of upper extremity paralysis, FES systems are often controlled by residual functional musculature to cycle between and activate grips. One study investigated how well three people with tetraplegia could use their shoulder position, wrist position, or wrist myoelectricity to control FES for opening and closing a grasp^11^, establishing the control systems used in many follow-up studies^6,9,12,13^. While successfully restoring functional grasps to paralyzed hands in these studies, the controllers are not intuitive beyond a single degree of freedom and would require extensive training for efficient usage. Furthermore, they are not generalizable solutions and people with severe tetraplegia, such as high cervical spinal cord injury, may not have sufficient residual function for such EMG or positional detectors.

Brain-machine interfaces (BMIs) may provide a more intuitive control system for upper extremity FES neuroprostheses that remains relevant to tetraplegia of more etiologies. By directly extracting intention information from the brain, BMIs have allowed people with paralysis to control and perceive a variety of end effectors, including controlling computers^14–17^, controlling multi-dimensional robotic arms^18,19^, and perceiving sensations^20–22^. As indicated by most tetraplegic participants of a survey, “brain-computer interface (BCI) control of [FES for hand grasp] would be ‘very helpful.’”^23^

Those who have translated BMIs to use with FES have successfully shown restoration of function in paralysis with monkeys^24–27^ and people^28,29^. For upper extremity restoration, brain-controlled functional electrical stimulation typically comes in discrete or continuous control forms. For example, Bouton et al. classified the user’s intentions with a support vector machine and delivered surface stimulation patterns to generate intended hand postures. For continuous control, Moritz et al. and Ethier et al. predicted and restored muscular force outputs as a continuous function of the spiking activities of few (Moritz et al.) or hundreds (Ethier et al.) of units. Badi et al. used spiking activities of tens of units to continuously control stimulation to just the temporarily paralyzed radial nerve in an object grasping task. Ajiboye et al. used a similar control system. They recorded units from hundreds of electrodes to continuously predict joint angles via optimal linear estimation. Then, those predicted joint angles were translated to stimulation patterns that generated the user’s intended movements. These studies showed clear improvements in abilities during paralysis, however, restoration of hand function only came in the form of discrete grasps. Even for regression-based algorithms, the tasks involved swapping between a hand opened or closed state. While undeniably important for completing common activities of daily living, there is insufficient understanding of how precisely FES can continuously control movements of the hand.

In this work, we leveraged our high-performance finger brain-machine interface^30–33^ to restore continuous movement to temporarily paralyzed prehensile fingers in a nonhuman primate (NHP) using functional electrical stimulation, the first of such a demonstration to our knowledge. When using the system, the monkey acquired virtual finger targets at a rate substantially higher than he could with his temporarily paralyzed hand. With two monkeys in an anesthetized state, we used target information to control FES directly and found that hysteresis, failure of muscular recruitment, and feedback latency substantially reduce target acquisition speed and create challenges for a BCFES neuroprosthesis user. Finally, we demonstrate, for the first time, that the simultaneous and independent movements of multiple finger groups can be controlled in a virtual environment by a brain-machine interface even when the monkey’s hand is temporarily paralyzed. Despite the performance reduction that was likely a result of the absence of sensory feedback, recalibrating the brain-machine interface using recalibrated feedback intention-training (ReFIT) restored performance to levels achieved by the brain-machine interface prior to temporary paralysis.

## RESULTS

### Restoration of Continuous Hand Function with Brain-Controlled Functional Electrical Stimulation in Temporary Paralysis

We first sought to estimate a nonhuman primate’s capability of continuously controlling his own temporarily paralyzed hand using a brain-controlled functional electrical stimulation neuroprosthesis. To do so, we trained a nonhuman primate to perform a finger task in which he controlled a virtual hand model using proportional movements of his physical fingers. A virtual target of a size equivalent to 15% or 16.5% of the range of motion (depending on the stage of BCFES training) was presented for movements of all four prehensile fingers together, and the monkey was first required to move his physical fingers such that the virtual fingers entered the target and held it continuously for 750ms. Then, we trained and tested a ReFIT Kalman filter (RKF) to predict the primate’s intended finger movements from spiking band power (SBP) recorded from Utah microelectrode arrays implanted in the motor-related hand area of precentral gyrus in real-time (see Methods)^32^. We used SBP, computed by taking the absolute value of the 300-1,000Hz band and averaging in 32ms bins, as we have previously shown it can extract firing patterns at lower signal-to-noise ratios, is more specific to the spiking rates of single units, and achieves as good or better prediction performance than threshold crossing rates^30^.

To demonstrate that we could control finger movements continuously with FES, we also implanted chronic bipolar stimulating electrodes into finger-related muscles of the forearm (see Methods for a list of electrode locations and quantities). Then, we delivered a lidocaine with epinephrine solution to surround the median, radial, and ulnar nerves just proximal to the elbow to temporarily block voluntary muscle contractions of the hand. We delivered electrical stimulation using the Networked Neuroprosthesis Evaluation System (NNP)^9^, which is a benchtop version of a human implantable device. We used the NNP to control the aperture of the hand’s fingers with stimulation patterns, or a number between 0% and 100% controlling the stimulation parameters delivered to all electrodes^28,34^. Then, we tested the monkey’s ability to control the virtual hand using his physical hand movements after the nerve block (purple route in Fig. 1a), using the RKF’s predictions to control the virtual hand directly (blue route in Fig. 1a), and using the RKF to control the stimulation delivered by the NNP with the virtual hand controlled directly by his physical hand movements (red route in Fig. 1a).

**Fig. 1.**
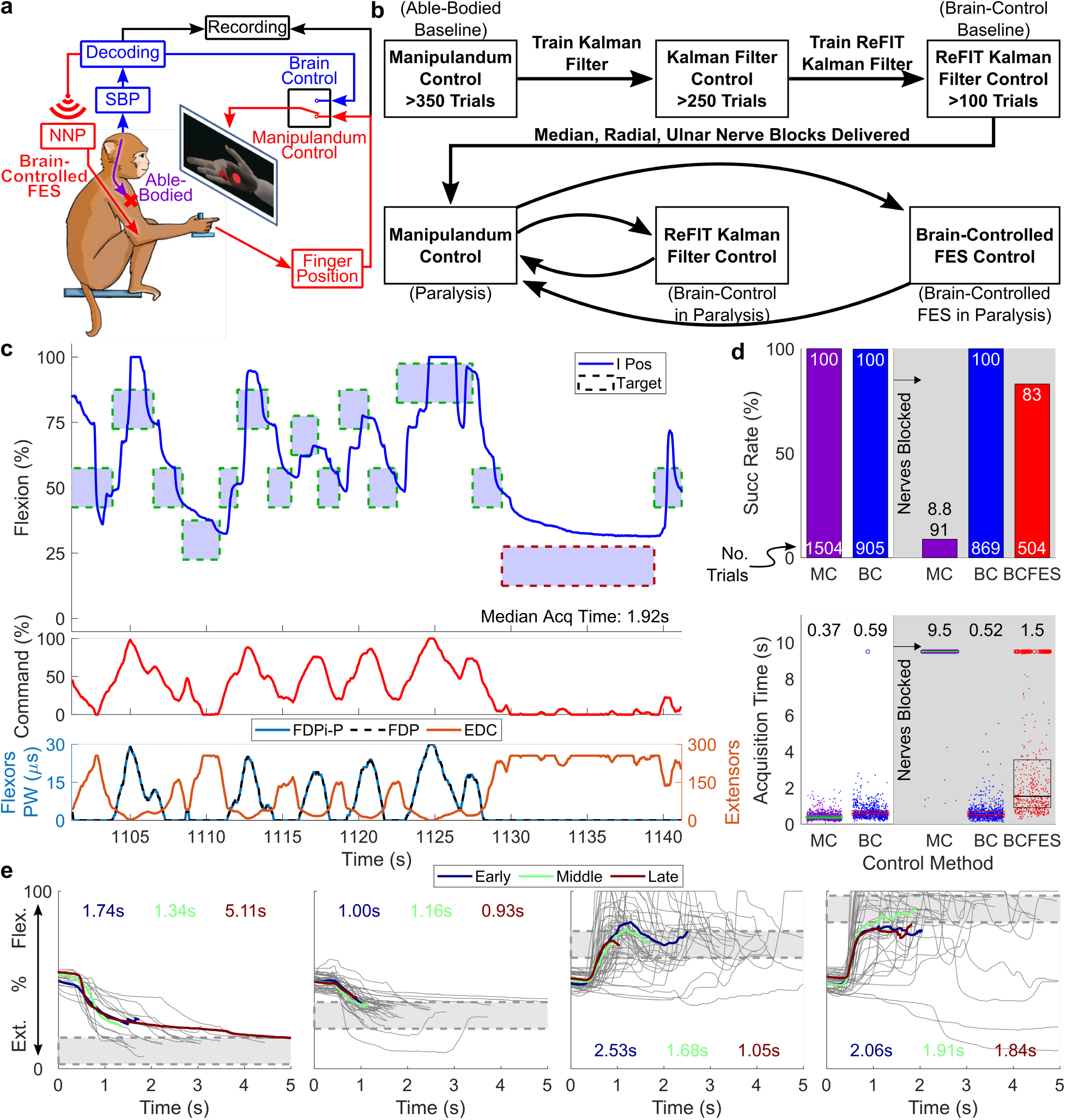
Brain-controlled functional electrical stimulation restores continuous hand function following regional blockade. (**a**) Diagram portraying the experimental control options. Purple represents able bodied control. This mode can be switched off via nerve block to the median, radial, and ulnar nerves. Blue represents brain control. Red represents brain-controlled functional electrical stimulation (BCFES). (**b**) Flow diagram of experimental procedures for each experimental day (four total days). (**c**) Example control traces under BCFES with stimulation command and delivered stimulation pulse widths. Green-border targets were successfully acquired, and red-border targets failed to trial timeout. I corresponds to index position. The targets were positioned in the active range of motion of the index finger, and all fingers moved as a group. (**d**) Target acquisition statistics for each method of control across all four experiment days. Boxes represent the 25^th^ and 75^th^ percentiles, with the midway line representing the median. Dots represent successful trials, and circles represent trials failed to timeout. MC corresponds to manipulandum control. BC corresponds to ReFIT Kalman filter (RKF) control. (**e**) All BCFES successful trial traces split by closest and furthest outer targets across all four experiment days. The blue, green, and red colored traces represent mean traces during early, middle, and late trial epochs, respectively. Median acquisition times for each target are displayed and colored according to the epoch they represent.

Fig. 1d shows single-trial statistics from all four experiment days under all control methods: manipulandum and RKF control prior to nerve block and manipulandum, RKF, and BCFES control following the nerve block. Using the BCFES system, Monkey N could acquire 83% of the presented targets (504 trials) in a median acquisition time of 1.5s. BCFES substantially improved control capabilities over Monkey N’s native hand movements following the nerve block, which had an 8.8% success rate (91 trials) and a median acquisition time of 9.5s, which is the trial timeout. Supplementary Video 1 portrays this comparison between BCFES and able-bodied control during the nerve block. Following the nerve block, Monkey N’s usage of the ReFIT Kalman filter to control the animated fingers did not change substantially, maintaining a 100% success rate with a median acquisition time of 0.52s (869 trials).

Fig. 1c shows a representative BCFES control example with decoded commands and delivered stimulus pulse widths included. Most failures of the BCFES were towards further extension targets, as Monkey N’s extensors fatigued quickly during BCFES usage. This is clearly showcased in Fig. 1e, which displays individual successful trial traces to each of the closest and furthest outer targets with early, middle, and late trials averaged into three highlighted traces of median length. Late trials to extension targets generally had lower success rates than during early trials, with only 1 successful acquisition of 13 far extension targets during the late epochs compared to 10 of 16 successes to the same target during early epochs. Recruitment of Monkey N’s EDC muscle was challenging even at maximum stimulation, which resulted in rapid fatiguing and his reliance on gradual recruitment to hit extension targets. Surprisingly, success rates remained high for flexion targets through all epochs and median acquisition times generally decreased in later trials (significance between late and early epochs for the nearest flexion target, *p* < 0.01; all others not significant, *p* > 0.01, two-tailed two-sample Wilcoxon rank sum test). This may be a result of Monkey N learning to better use the BCFES system or muscle fatigue improving the controllability of the stimulation.

### High-Speed Time to Target with Target-Controlled FES

We next sought to better understand how well our FES solution worked as a standalone finger prosthesis without BMI control. Such an analysis can be informative about what challenges a user must consciously balance when using a BCFES neuroprosthesis. To investigate this, we anesthetized Monkey N and Monkey W (who was implanted with acute FES electrodes, see Methods) with propofol and used target information to control the stimulation command on two experiment days each. Supplementary Fig. 1 illustrates the simple proportional controller, where at the beginning of a trial, the stimulation command is set to the target position (in the range of 0% to 100%). Then, to provide any necessary minor corrections ideally without overshooting the target, the stimulation command was increased or decreased by approximately 1%, near the minimum control resolution, every iteration depending on the direction of the target from the current position of the finger.

Fig. 2a and b illustrates acquisition time statistics for all trials with target-controlled stimulation for both monkeys, with colors indicating the direction of movement in the center-out task. Overall, times to target were quick for Monkey N (median 0.86s) and slower but still rapid for Monkey W (median 1.7s), comparable to typical times to target during BMI use. Times to target for flexions were generally faster with Monkey N (median 0.73s) than extensions (median 1.4s). This was likely due to Monkey N’s flexor muscles showing better recruitment than the extensors which quickly exhibited muscle fatigue. This fatigue also resulted in substantially higher variability in times to target when reaching towards extension targets than flexion or central targets.

**Fig. 2.**
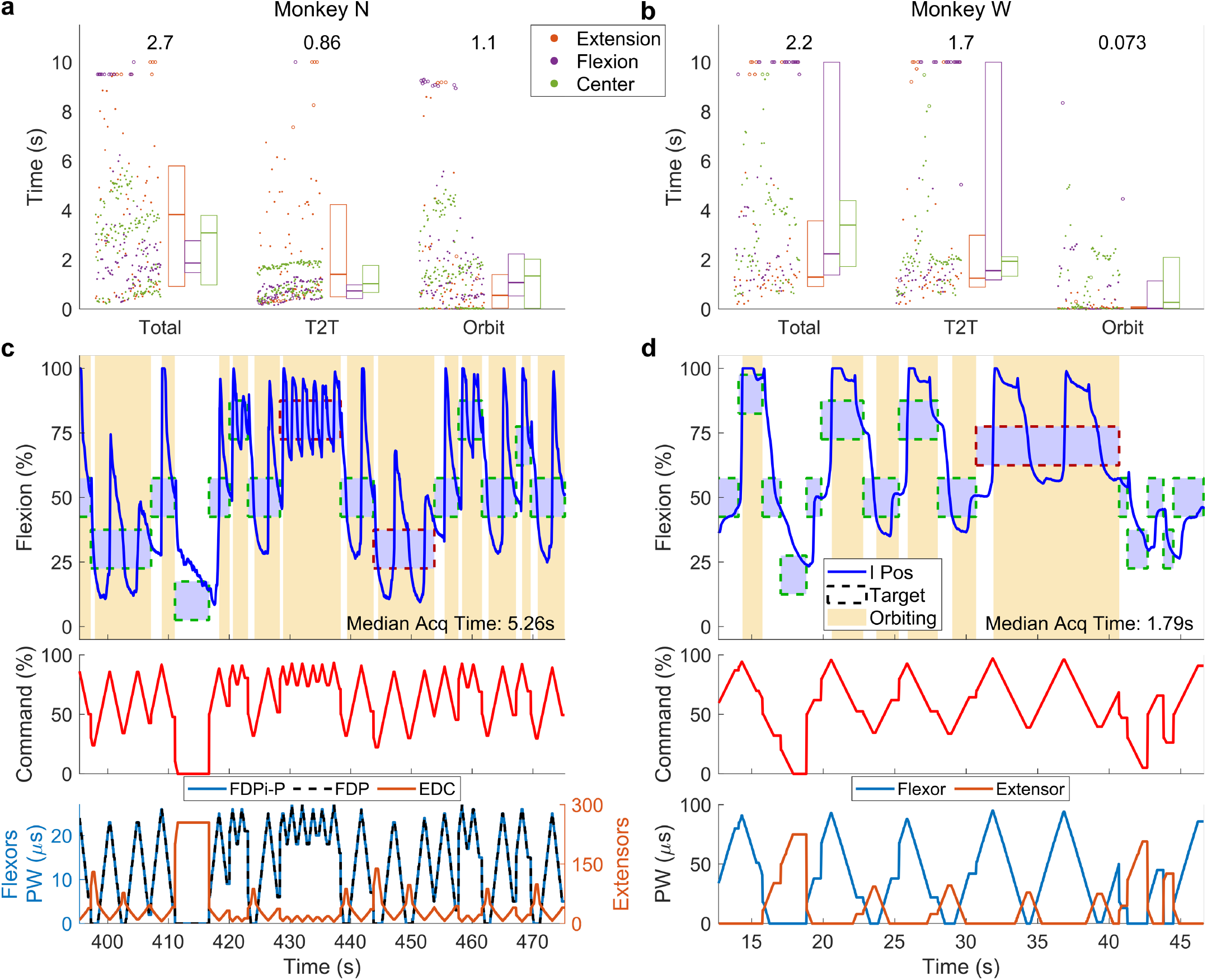
Target-controlled functional electrical stimulation achieves high-speed time to target. (**a**) Target acquisition statistics for target-controlled stimulation with Monkey N for all two experiment days. T2T means the time to target, or the amount of time to first touch the target. OT means the orbiting time, or the amount of time between first touching the target and successfully acquiring it, less the required hold time. Boxes to the right represent the 25^th^ and 75^th^ percentiles, with the midway line representing the median for each type of target. Dots represent successful trials, and circles represent trials failed to timeout. (**b**) Same as **a** but for Monkey W’s two days of experiments. (**c**) Example control traces under target controlled stimulation for Monkey N with stimulation command and delivered stimulation pulse widths displayed in parallel. Green-border targets were successfully acquired, and red-border targets failed to trial timeout. Yellow regions represent periods of orbiting the target. I means index position, for which the target was presented despite the whole hand being controlled by stimulation. **(d**) Same as **c** but for Monkey W.

Interestingly, orbiting, or the action of dialing into a target after first hitting it, was a challenge in both monkeys (median 1.1s for Monkey N, median 0.073s for Monkey W). We expected FES to slowly recruit muscles with increasing stimulus and not activate them as well as it did. The individual trial traces in Fig. 3 and the top plots in Fig. 2c and d illustrate the orbiting, where the yellow regions show that orbiting occupies the majority of the time axis. Hysteresis in muscular activation from stimulation, particularly for the flexors, caused many of these orbiting periods, evidenced by the frequent passing of the target from both sides. In contrast, gradual recruitment of the extensors helped reduce orbiting (for extension vs. flexion and center targets: median 0.49s vs. 1.24s in Monkey N, *p* = 5.6 × 10^−3^; 0.040s vs. 0.080s in Monkey W, *p* = 8.8 × 10^−3^, two-tailed two-sample Wilcoxon rank sum test). Notably, the acute electrodes implanted in Monkey W yielded a drastically lower median orbiting time, particularly during small bursts of consecutive trials without flexion targets as seen in Fig. 2d and Fig. 3. We hypothesize these acute electrodes enabled smoother recruitment of the muscles, potentially due to the lack of scar tissue surrounding the electrode sites. Overall, orbiting poses a substantial challenge to usage of a continuous FES system, causing 16 of 20 failures in Monkey N and 6 of 28 failures in Monkey W.

**Fig. 3.**
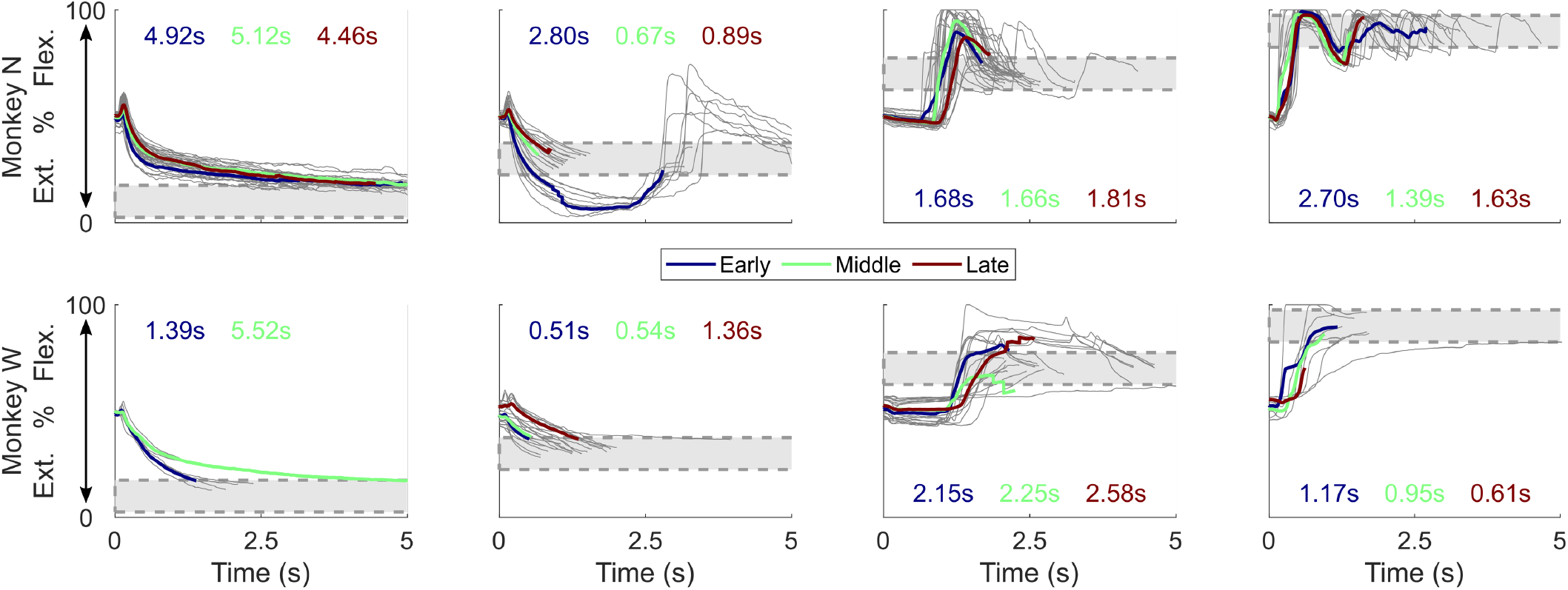
Single trial traces for target-controlled functional electrical stimulation. All target-controlled FES trials split by closest and furthest outer targets. The blue, green, and red colored traces represent mean traces during early, middle, and late trial epochs, respectively. Median acquisition times for each target are displayed and colored according to the epoch they represent.

### High Performance Two-Finger Brain-Control in Temporary Paralysis

To close the gap towards a multi-finger brain-controlled functional electrical stimulation neuroprosthesis, we next wanted to investigate how well our brain-machine interface systems could predict multi-finger movements following nerve block, i.e. controlling animated fingers when the hand has been temporarily paralyzed. Similar to our one-dimensional finger task discussed thus far, we trained a two-dimensional RKF to predict the simultaneous and independent movements of two finger groups (the index finger separate from the middle, ring, and small fingers as a group) prior to blocking the nerves^31^. Then, we used the RKF to predict the two-dimensional finger movements in real-time following the nerve block on two separate days.

Fig. 4 illustrates the results of these experiments, where prior to the nerve block, the monkey’s capabilities of completing the task with his physical hand and the RKF were high (100% success rate for both and 0.39s and 0.75s median acquisition times, respectively). Following the nerve block, the monkey’s acquisition rate when using his physical hand dropped drastically (2.5% success rate due to chance acquisition, with 9.5s median acquisition time), as expected due to the inability to physically move his fingers.

**Fig. 4.**
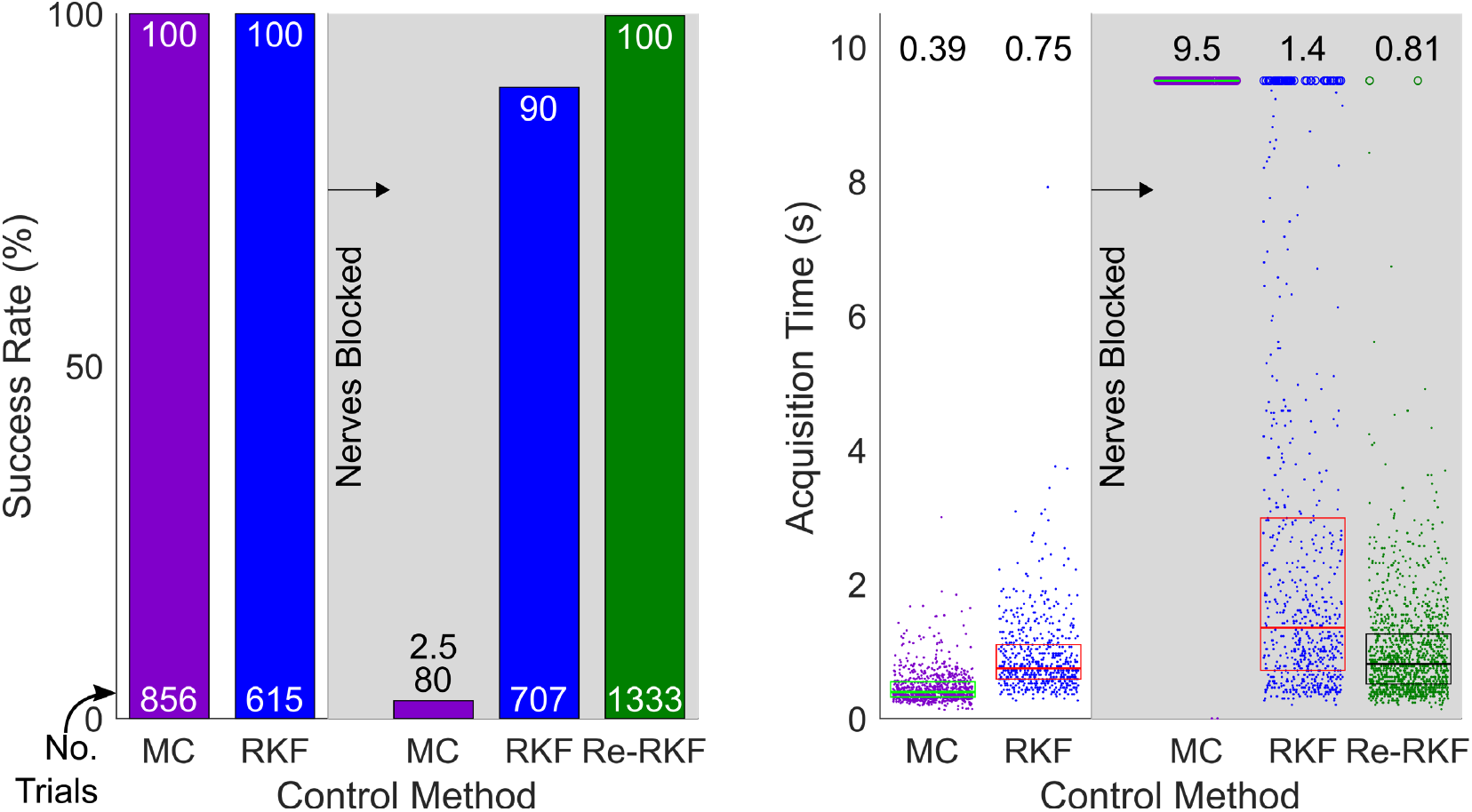
Recalibrated RKF restores high-performance control of virtual multi-finger movements following nerve block. Success rates and acquisition times for manipulandum, RKF, and recalibrated RKF (Re-RKF) two-finger control before and after nerve blocks are shown. Numbers shown atop the acquisition times plot represent the median acquisition time. Boxes within the plots show median in the center with the edges representing 25^th^ to 75^th^ percentiles. RKF and Re-RKF acquisition times were not statistically significantly different (*p* = 0.67, two-tailed Wilcoxon rank sum test), but all other acquisition time comparisons were statistically significant (*p* < 0.001, two-tailed Wilcoxon rank sum test).

Then, we transitioned back to investigating usage of the BMI following the nerve block. When testing the RKF, we surprisingly found a substantial drop in performance (90% success rate with 1.4s median acquisition time) despite no direct cortical interventions. In an attempt to restore performance, we performed the ReFIT training procedure an additional time using the closed-loop RKF trials Monkey N performed following the nerve block. We termed this second-stage RKF the Re-RKF, indicated in green in Fig. 4. After recalibrating, closed-loop control with the Re-RKF returned to the level of performance achieved by the RKF prior to the nerve block (100% success rate for both and 0.75s vs. 0.81s median target acquisition time between RKF prior to the nerve block and Re-RKF, respectively; *p* = 0.67, two-tailed Wilcoxon rank sum test).

The abrupt drop in BMI performance following the nerve block motivated us to investigate what may have been its cause. When we blocked median, radial, and ulnar nerves, not only were the descending motor commands prevented from reaching their muscular targets, but we think proprioceptive and sensory feedback were also partially or completely blocked from transduction back to cortex. The RKF trained on able-bodied behavior might have learned to depend on some of this sensory information, which then becomes absent following a nerve block, possibly accounting for the performance drop. Fortunately, during these experiments, we simultaneously recorded sensory activity from a separate Utah array implanted in sensory-related postcentral gyrus alongside the motor-related precentral gyrus array used for closed-loop control. This gives us an avenue through which we can investigate how the nerve block impacts BMI control with the RKF.

We first asked whether we usually had sensory tuning to the task’s kinematics represented on an array in sensory-related postcentral gyrus. To do so, we used historical data from Utah arrays implanted into the postcentral gyri of two animals (Monkey W and a previous implant in Monkey N). We trained an open-loop, offline Kalman filter to use SBP recorded from postcentral gyrus to predict the one-finger movements measured by the manipulandum, doing so with intra-class correlations (ICCs) of 0.81 and 0.58 for Monkeys N and W, respectively (Fig. 5a). This suggests that the sensory data usually has good linear tuning to finger kinematics, which a regression would tend to find. We also saw the same result for the two finger task with Monkey N’s active postcentral gyrus array, as shown by the purple bars in Fig. 5d (mean 0.65 ICC across both experiment days).

**Fig. 5.**
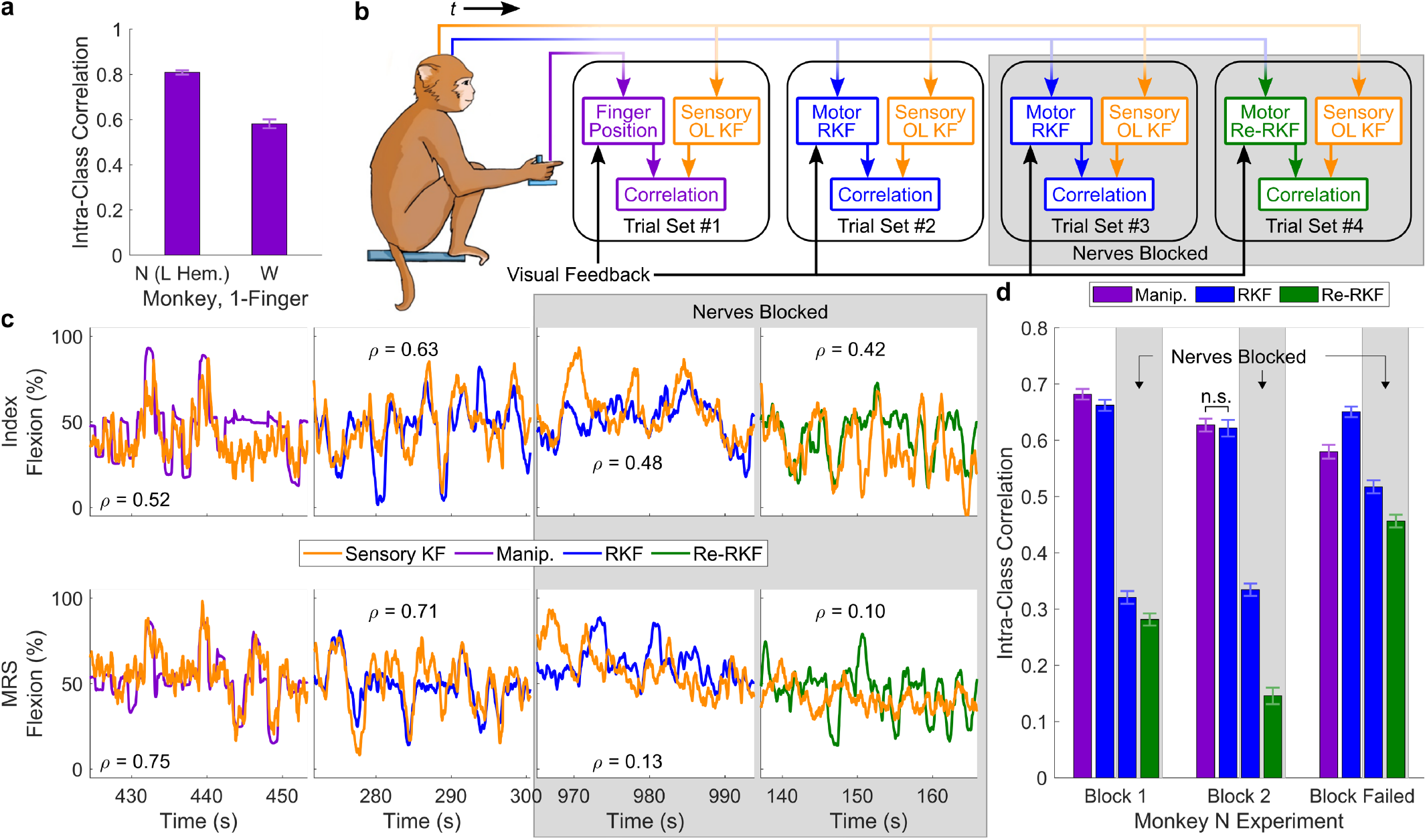
Absence of sensory feedback may cause closed-loop performance reduction following nerve block. (**a**) SBP in sensory-related postcentral gyrus accurately predicts one-finger movements measured with the manipulandum in two monkeys. (**b**) Description of analysis. SBP was recorded from Monkey N’s postcentral gyrus implant synchronously with manipulandum measurements of two-finger movements and the SBP from Monkey N’s motor-related precentral gyrus implant. For each comparison, we trained an open-loop Kalman filter to predict the virtual control outputs (manipulandum, RKF, or Re-RKF) from postcentral gyrus SBP recorded at the same time. The predictions from the sensory Kalman filter were intra-class correlated with the control outputs during each set of trials. (**c**) Example cross-validated open-loop position predictions from the sensory Kalman filter (orange) overlaid on the control outputs. The horizontal time axis represents the seconds into each usage instance of each closed-loop control method. Index and MRS were controlled simultaneously by the monkey with traces split here for clarity. The included correlations represent each snippet’s intra-class correlation. (**d**) Intra-class correlation coefficients between the two-finger sensory Kalman filter predictions and each control method, with positions combined across fingers and all collected data from each closed-loop control method. Error bars represent the upper and lower bounds with an alpha level of 0.001. Data represent experiments with nerve blocks on two separate days with one additional day in which the nerve block failed (85% success rate with manipulandum control post-block) to show maintained sensory prediction of closed-loop RKF and Re-RKF predictions. All comparisons within each day were significantly different (*p* < 0.001, one-sided two-sample *z*-test on the Fisher-transformed intra-class correlation coefficients), except those labelled with n.s.

Next, we wondered if sensory tuning remained representative of the kinematics predicted by the BMI, both before and after the nerve block (illustrated by Fig. 5b with results summarized in Fig. 5c and d). Prior to the nerve block, an offline sensory Kalman filter accurately predicted the closed-loop RKF predictions, but performance drastically dropped following the nerve block and during Re-RKF use (mean 0.64, 0.33, then 0.21 ICC, respectively, *p* < 0.001 between all comparisons, one-tailed two-sample *z*-test on the Fisher-transformed ICC). This suggests that there was a loss of information transfer from sensory-related to motor-related cortical areas during the finger task without sensory feedback. Therefore, for the Re-RKF to restore closed-loop performance, performing the additional stage of ReFIT likely fit an observation model that was not dependent on sensory information.

## DISCUSSION

Modern FES has primarily focused on restoring discrete grasps and grips but the extent to which FES can continuously control end effectors is not well understood. In this work, we have demonstrated that an NHP can recover a substantial amount of continuous finger function following temporary paralysis of the hand by using a brain-controlled functional electrical stimulation system. The BCFES system allowed the monkey to dramatically exceed the performance of his native hand following targeted regional blockade of the median, radial, and ulnar nerves, though it could not achieve the levels of performance of the able-bodied hand prior to temporary paralysis nor the same level of performance as when controlling an animation. By using information about the presented target to control stimulation without voluntary control from the monkey, we found that hysteresis, muscle recruitment failure, and controller latency may be the primary challenges in using a BCFES prosthesis. Fortunately, these challenges can be partially mitigated by the subject via brain-control. Finally, we have also presented the first demonstration that the simultaneous and independent movements of multiple finger groups can be continuously predicted with high performance when the hand is temporarily paralyzed despite the absence of proprioceptive and sensory feedback. Recalibrating a ReFIT Kalman filter restored performance of the two-finger decoder to equivalent levels as the original ReFIT Kalman filter used prior to the regional block. This is a particularly surprising result, as it suggests that brain-controlled FES performance in a paretic scenario may not be limited by the BMI decoder but rather by current FES technologies.

Interestingly, our analysis of sensory representation in motor cortex following the nerve block suggests that the loss of sensory feedback may have been the cause for the drop in performance. When using sensory-related postcentral gyrus activity to predict the closed-loop RKF and Re-RKF predictions following induction of nerve block, similar to analyses done by others^35^, we noticed a substantial drop in intra-class correlation compared to before blockade. We hypothesize that some of the precentral gyrus SBP channels that the RKF found valuable contained some information transferred from postcentral gyrus, which is not particularly surprising given that sensory-related postcentral gyrus accurately represents hand joint angles^36^ and projects to and receives from motor-related precentral gyrus^37–39^. Since precentral gyrus did not have any sensory inputs during usage of the RKF following the nerve block, training the Re-RKF decoder used purely motor-related activity, which we think is what allowed it to function with high performance in the absence of sensory feedback. Although this hypothetical explanation is intriguing, further experimental validation will be necessary to support it, particularly without the capability of verifying the complete loss of sensation with an animal model.

When investigating the capability of FES to continuously control finger movements without voluntary intervention from the monkey, we found substantial orbiting to reach flexion and extension targets that was likely an outcome of hysteresis, failure of muscle recruitment, and general fatiguing^40–42^. Latency in between a command and the behavioral response resulting from stimulation could also contribute to oscillation about a target, where we found in Supplementary Fig. 5 that the latency with BCFES was substantially higher than the latency with the RKF and manipulandum control systems. Therefore, this latency was also likely an obstacle for Monkey N during closed-loop BCFES, though learning to use the system as shown in Fig. 1 enabled him to reduce these oscillations. Additional feedback controllers, such as a well-tuned proportional-integral-derivative controller^43^ or a cosine similarity controller^44^, could reduce the cognitive load on the user when combating these systematic challenges of BCFES. The error between the decoder’s predictions and an estimate of the present positions of the fingers, which could be captured by a glove worn by the user, could inform the error signal to the controller.

The BCFES results presented here expand upon the BCFES accomplishments of others^24–26,28,29^ by beginning to investigate the realm of continuous control with FES, using similar technologies as what was previously used in clinical trials. Although our monkey model of temporary paralysis should not be considered equivalent to chronic tetraplegia, one could consider the model to be representative of an ideal case of tetraplegia without typical pathologies, such as spasticity or contractures. Even though the results presented here might represent best-case scenarios given the techniques and methods used, a user may be able to perform more natural hand movements and use the device for a wider variety of tasks just by adding continuous control to grasp and force. Furthermore, combining these techniques with sensory feedback via intracortical microstimulation^20,21^ may enhance control capabilities. Although we have shown sensory feedback is not required for cortical control of dexterous finger movements, it may help with the accuracy of BCFES movements as it has been shown to improve motor control^20^. Including a hand-state measurement glove, as discussed in the previous paragraph, could acquire the necessary signals to deliver valuable proprioceptive and force-related feedback in addition to improving BCFES performance.

For the purposes of translating BCFES for functional restoration beyond laboratory use, the technologies presented in this manuscript all represent low-power solutions end-to-end. The stimulation system used in this work is an evaluation system for the Networked Neuroprosthesis, a low-power, wireless, completely implantable stimulator presently undergoing a clinical trial (NCT02329652). The brain-machine interface used the 300-1,000Hz spiking band power, a low-power neural feature that is specific to single-unit activity despite a ten-fold reduction in processing requirements. We have already demonstrated SBP’s efficacy on embedded and integrated systems^45–49^, and others have used it in similar brain-machine interface and BCFES applications^28,50,51^. While much work remains in system integration and safety validation, BCFES has a strong potential for full-time use in people with paralysis.

## METHODS

All procedures were approved by the University of Michigan Institutional Animal Care and Use Committee.

### Implants

We implanted two adult male rhesus macaques with Utah microelectrode arrays (Blackrock Microsystems, Salt Lake City, UT, USA) in the hand area of pre- and postcentral gyri, as described previously^31–33^. Monkey N was first implanted with two 64-channel arrays in left hemisphere precentral gyrus and one 96-channel array in left hemisphere postcentral gyrus. Data from the postcentral gyrus array was only used to investigate how well sensory information could predict one finger movements measured by a manipulandum. Monkey N was age 5 years and was 37 days post implant for this data collection. Later, connections to these arrays were cut and Monkey N was implanted with two new 64-channel arrays in right hemisphere precentral gyrus and one 96-channel array in right hemisphere postcentral gyrus. Monkey N was age 9 and between 867 and 1,071 days post cortical implant for all other data collected in this study, which resulted from these arrays. Monkey W was implanted with one 96-channel array in each of left hemisphere precentral and postcentral gyri. Monkey W was age 6 years and was 590 days post implant at the time of data collection. All of Monkey N’s right hemisphere arrays, only Monkey N’s left hemisphere postcentral gyrus array, and only Monkey W’s postcentral gyrus array were used in this study. Pictures of all implants can be found in Supplementary Fig. 2.

In a separate surgery, we implanted Monkey N with 86cm chronic bipolar intramuscular electromyography recording electrodes (similar to PermaLoc™ electrodes, Synapse Biomedical, Inc., Oberlin, OH, USA). These electrodes were limited to 5μC of charge per pulse. Electrodes were implanted as described previously^31^. Briefly, a single radial-volar incision was used to access flexor muscles, and electrodes were placed in flexor digitorum profundus-index (1x near the nerve entry point, 1x distal near the wrist), flexor digitorum profundus-MRS (1x), flexor pollicis longus (1x, unused in this study), flexor carpi radialis (1x, unused in this study), and flexor carpi ulnaris (1x, unused in this study). Electrodes were secured intramuscularly using non-absorbable monofilament suture. After closing the radial-volar incision, a single dorsal-ulnar incision was used to access the extensors. Electrodes were placed in extensor digitorum communis (1x), extensor indicis proprius (1x, unused in this study), extensor carpi radialis brevis (1x, unused in this study), and extensor pollicis longus (1x, unused in this study). Electrodes were tunneled proximally to an interscapular exit site and connected to the standard PermaLoc™ connector. All incisions were closed in a layered fashion using absorbable sutures and leads were stitched to the exit site. Following implantation, the monkey persistently wore a Primate jacket (Lomir Biomedical, Inc., Malone, NY, USA). Monkey N was between 195 and 399 days post FES electrode implant for all data collected in this study.

To investigate continuously-controlled FES in Monkey W, we acutely implanted bipolar fine wire electrodes (019-475400, Natus Medical Inc., Middleton, WI, USA) in his left forearm. After induction of anesthesia with propofol (see methods below), we targeted flexor digitorum profundus-MRS and extensor digitorum communis anatomically and observationally by manually flexing and extending the fingers.

### Feature Extraction

All processing was done in MATLAB versions 2012b or 2019b (Mathworks, Natick, MA, USA), except where noted.

Precentral gyrus (motor-related) SBP was acquired in real-time during the experiments (see the subsequent section for a description of data flow). To do so, we configured the Cerebus neural signal processor (Blackrock Microsystems) to band-pass filter the raw signals to 300-1,000Hz using the Digital Filter Editor feature included in the Central Software Suite version 6.5.4 (Blackrock Microsystems), then sampled at 2kSps for SBP. The continuous data was streamed to a computer running xPC Target version 2012b (Mathworks), which took the magnitude of the incoming data, summed all magnitudes acquired in each 1ms iteration, and stored the 1ms sums as well as the quantity of samples received each 1ms synchronized with all other real-time experimental information. This allowed offline and online binning of the neural activity to create larger bin sizes, such as the 32ms used in this work, with 1ms precision. We masked channels for closed-loop decoding to those that were not saturated with noise and had contained morphological spikes during the experiment or at some time in the past, as SBP could possibly extract firing rates of low signal-to-noise ratio units remaining represented on such channels^30^.

Postcentral gyrus (sensory-related) SBP was recorded to disk for later offline synchronization. The pedestals for the arrays implanted in postcentral gyrus were connected to a CerePlex Direct (Blackrock Microsystems) via a CerePlex E (Blackrock Microsystems), which either collected the raw data at 30kSps or band-pass filtered the incoming signals to 300-1,000Hz using the Digital Filter Editor feature, then sampled at 2kSps for SBP. To synchronize the postcentral gyrus SBP activity after the experiment, we used the Sync Pulse functionality included in Central. The unique Sync Pulses were recorded by both the Cerebus and the CerePlex direct, enabling synchronization of the recordings from both systems offline. Then, postcentral gyrus SBP was filtered to 300-1,000Hz if not previously done, absolute valued, and accumulated in equivalent windows as the precentral gyrus SBP used for comparison using MATLAB R2019b.

### Experimental Setup

The experimental apparatus used for these experiments is similar to what was described previously^31–33^. Briefly, the monkeys’ Utah arrays were connected to the patient cable (Blackrock Microsystems) and neural data (as described previously) were streamed to the xPC Target computer in real-time via a User Datagram Protocol packet structure. The xPC Target computer coordinated several components of the experiments, including coordinating target presentation, acquiring measured finger group positions from one flex sensor per group (FS-L-0073-103-ST, Spectra Symbol, Salt Lake City, UT, USA), and transmitting finger positions along with target locations to an additional computer simulating movements of a virtual monkey hand (MusculoSkeletal Modeling Software; Davoodi et al., 2007). Task parameters, states, and neural features were stored in real-time for later offline analysis.

One additional functionality was implemented in the xPC Target computer to facilitate real-time FES control. We used an RS-232 interface to the Networked Neuroprosthesis Access Point that comprised of an MSP-EXP430F5529LP evaluation board (Texas Instruments Inc., Dallas, TX, USA), an Evaluation Module Adapter Board (Texas Instruments), a CC1101EMK433 evaluation kit (Texas Instruments), and a MAX3222E RS-232 level shifter (Maxim Integrated, San Jose, CA, USA). Stimulation commands for one pattern were transmitted at 115,200 baud, which were sent wirelessly to the NNP power module to be configured into pulses delivered (see Functional Electrical Stimulation and Patterns sections below).

### Behavioral Task

We trained Monkeys N and W to acquire virtual targets with virtual fingers by moving his physical fingers in a one- or two-finger task, similar to what we have previously published^31,33^. During all sessions, the monkeys sat in a shielded chamber with their arms fixed at their sides flexed at 90 degrees at the elbow, resting on a table. The monkeys had their left or right hand (contralateral to cortical implants and ipsilateral to intramuscular implants) placed in the manipulandum described previously^31^. During manipulandum control (not FES), Monkey W had the flexion measuring sensors (FS-L-0073-103-ST, Spectra Symbol Corp., Salt Lake City, UT, USA) taped directly to his index finger, but could move all fingers freely. During historical data collection from Monkey N’s left hemisphere (for the sensory analysis), the manipulandum degree of freedom for his MRS fingers was locked such that they were at full extension, while the index finger was free to move across its full range of motion. During all other Monkey N’s one-finger tasks in this study, the manipulandum’s degrees of freedom were locked together so that Monkey N could only move his fingers together. During Monkey N’s two-finger task, he was free to move both the index and MRS finger groups freely in their degrees of freedom within the manipulandum. The monkeys sat in front of a computer monitor displaying the virtual hand model and targets described previously.

Each trial began with one spherical target per finger degree of freedom appearing along the one-dimensional movement arc. Each target occupied 15% of the full arc of motion of the virtual fingers with two exceptions. BCFES trials with Monkey N during his first usage of the system had a 16.5% target size (94 of 424 presented trials) and all historical trials used to estimate how well postcentral gyrus could predict one-finger movements with Monkey W used a 14.25% target size. Targets were presented in a center-out pattern, every other target was presented at rest (halfway between full flexion and full extension or 50% as illustrated in figures), and the non-rest targets were randomly chosen between 20%, 30%, or 40% flexion or extension from rest. For a successful trial, the monkeys were required to move the virtual fingers into their respective targets and remain there for 750ms continuously in able-bodied manipulandum control or 500ms continuously in post-block manipulandum control, brain control, and BCFES modes. During historical trials with Monkey W estimating how well postcentral gyrus predicted one-finger movements during able-bodied manipulandum control, hold time was 500ms. If the monkeys could not acquire and hold the target within 10s, the trial was deemed unsuccessful and the target was placed at center repeatedly until successfully acquired. Upon successful target acquisition, the monkeys received a juice reward, which was modulated to maintain motivation levels.

### Nerve Block Procedure

Temporary paralysis of the hand was achieved by delivering a solution of lidocaine (2%) and epinephrine (1:100,000) intramuscularly at three peripheral nerves (radial, median, ulnar), in the upper arm just proximal to the elbow. The solution was delivered so that it completely surrounded each nerve. Supplementary Fig. 3 presents some example snapshots of the median, radial, and ulnar nerves from Monkey N under ultrasound. The lidocaine/epinephrine solution was either purchased premixed (NDC 0409-3182-11) or compounded using stock solutions of lidocaine (2%) and epinephrine 1 mg/mL. Delivery was done under ultrasound (Lumify L12-4 broadband linear array transducer, Philips Healthcare, Best, NL, paired with Samsung Galaxy Tab S6, Samsung Electronics, Suwon, KOR) using a 20 gauge echogenic needle (B.Braun #33642, B.Braun, Melsungen, DE). The NHP was placed in a restraint chair and the arm was manually restrained. In some experiments, prior to delivering the block, pure lidocaine (2%) was injected subcutaneously to the target injection sites to ease with comfort during the injection, and approximately 6-7mg/kg (4-6mL) of lidocaine in the lidocaine/epinephrine solution (less than that was ineffective) was used cumulatively for each blocking procedure. Block onset typically occurred after 20-30 minutes and lasted up to 2 hours. There were at least two days between blocking procedures on the same animal. To guarantee the nerve block was still active at any time during an experiment, we would occasionally disable the BCFES system and the decoder, allowing the monkey to complete trials with his native anatomy as capable.

### Propofol Anesthesia

To investigate the capabilities of functional electrical stimulation alone without voluntary activation from the monkey, we lightly anesthetized Monkeys N and W with propofol.

After placing the monkey in the primate chair, comfortably restraining his head, and comfortably restraining his arms, we placed an IV catheter in the cephalic vein just distal to the monkey’s right elbow, contralateral to the arm being used for FES experiments. We then flushed the catheter with sterile saline to ensure patency. We delivered a 2.5-3.0mg/kg bolus with additive 0.2mg/kg boluses of propofol until the primate was visibly unconscious to induce a light plane of anesthesia. Following induction, light anesthesia was maintained with a constant rate infusion of propofol at 7.5mg/kg/hr and supplemental boluses of 0.2mg/kg were given as needed to maintain the desired plane of anesthesia.

Oxygen supplementation was provided with a nasal cannula. Heart rate, respiratory rate, body temperature, and blood oxygen levels were monitored throughout the experiment with a pulse oximeter and pediatric ear thermometer. Blood oxygen levels never dropped below 90% and were always greater than or equal to 98% after providing supplemental oxygen.

Anesthetic depth was measured with jaw tension, respiratory rate, and heart rate. After a stable and light plane of anesthesia was reached, we proceeded with functional electrical stimulation. Anesthetic events/procedures occurred no more than once per week and for no longer than two hours with most lasting one hour.

### Functional Electrical Stimulation

We delivered stimulation using the Networked Neuroprosthesis Evaluation System^9^. Fig. 6 illustrates how the system was connected to Monkey N. The evaluation system contains the same circuitry used as implants in people (NCT02329652) but in a form factor more conducive of experimentation and debugging. Our evaluation system consisted of one power module, including three 1,000mAh batteries to power the system and deliver stimulation (PRT-13813, SparkFun Electronics, Niwot, CO, USA), and three 4-channel pulse generator modules. The system could deliver 12 total monopolar stimulation channels. All stimulation was current-controlled and delivered at 10mA with a 32ms inter-pulse interval. Pulses were charge-balanced biphasic with a square cathodic pulse delivered first and the subsequent anodic pulse exponentially balancing charge. Pulse delivery was staggered within each pulse generator module to avoid too large of a current draw than could be provided by the power module. The pulse for channel 1 of a given pulse generator module was delivered 20ms following the reception of the stimulation command (the time required to prepare the pulse based on the command), and each subsequent channel delivered its pulse 1ms after the prior channel. All pulse generator modules received the stimulation command simultaneously.

**Fig. 6.**
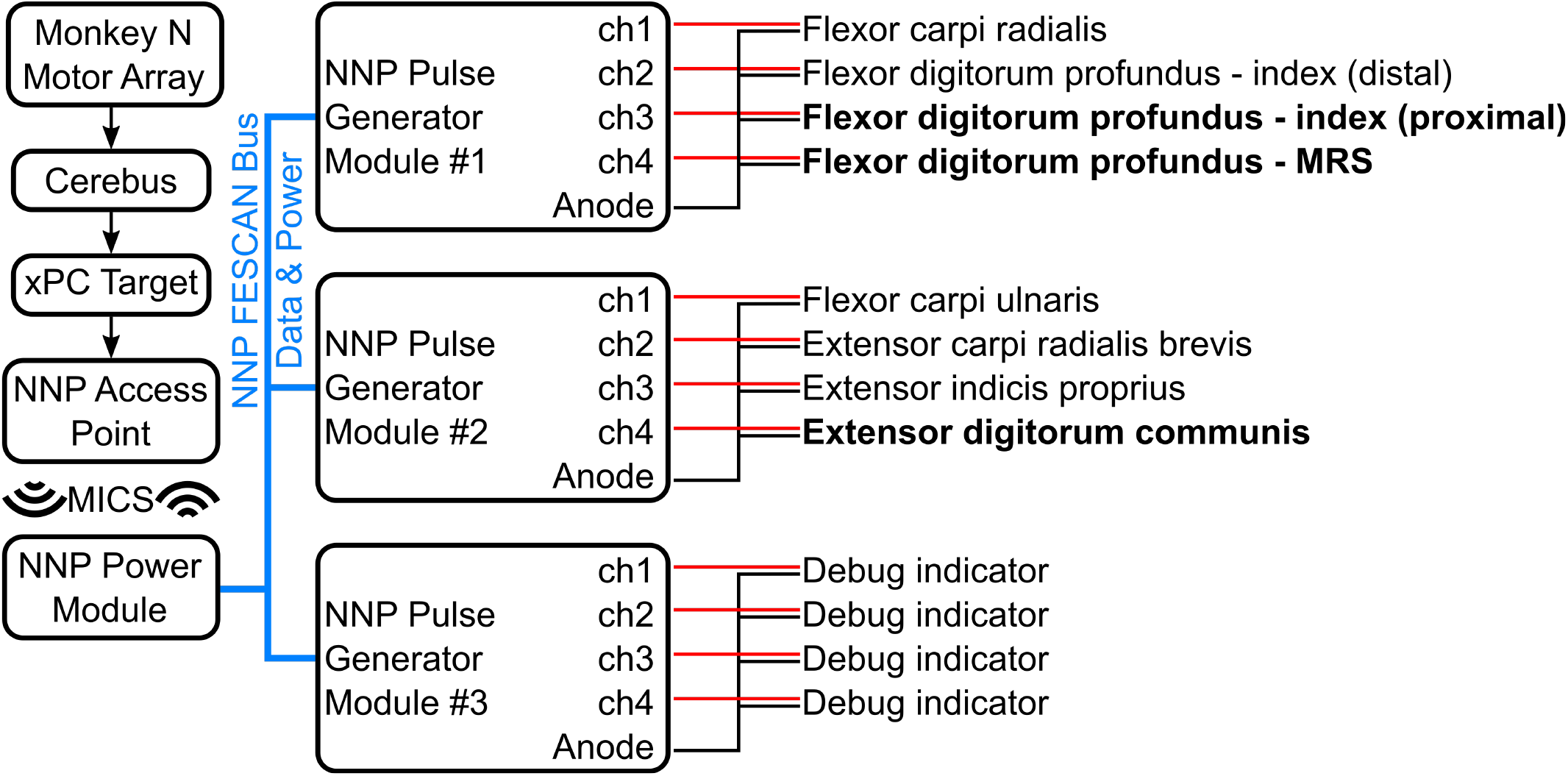
Connection diagram for Monkey N’s FES electrodes. Monkey W’s extensor and flexor were connected to NNP Pulse Generator Module #2 ch1 and ch2, respectively. Bolded muscle names represent muscles that were used in this study, though all muscles listed were connected to the system during testing. Each muscle has a red and black wire to represent the connections of each contact for each bipolar lead. Debug indicators were used to display system status during an experiment using light-emitting diodes and were not connected to the animal in any way. MICS refers to the Medical Implant Communication Service radio band that was used for wireless communication between the experimental system and the NNP, providing wireless isolation between the recording and stimulation systems (since the NNP was powered by batteries).

To adapt our bipolar leads to function with monopolar stimulation, we only used one electrode of each pair to deliver current. With Monkey N’s chronic electrodes, the electrode that was implanted further inside the muscle was used for monopolar stimulation. The second electrode of each bipolar pair that was further outside the muscle were tied with all other second electrodes connected to one pulse generator module. Each group of four electrodes that were further outside the muscle were connected to that pulse generator module’s current return. With Monkey W’s acute electrodes, we connected the NNP to each electrode pair using hook grabbers. One electrode of each pair was connected to one monopolar pulse generator, and one of the two remaining disconnected electrodes was connected to the current return of the NNP. Due to the variability in placement of the acute electrodes with Monkey W, occasionally a muscle would contract due to the returning current though the muscle. If this occurred, we would instead tie the current return to a surface electrode (NC0748095, Biopac Systems, Inc., Goleta, CA, USA) with conductive gel (SignaGel, Parker Labs, Inc., Fairfield, NJ, USA) as needed, placed either on the chest or back of the neck.

Prior to usage of the BCFES or target-controlled FES systems, the flex sensors were recalibrated to the range of motion that could be achieved independently by FES. During some BCFES and target-controlled FES experiments, small wooden stints were taped to the monkey’s fingers following the nerve block. FES would occasionally flex the distal and proximal interphalangeal joints within the manipulandum in a way that could not be measured by the manipulandum and prevent extension of the manipulandum. Without electrodes implanted in all hand muscles, including the intrinsics, FES could not independently re-extend the fingers. The stints helped with keeping the fingers straight and flexing primarily at the metacarpophalangeal joint. Additionally, during BCFES experiments, we would occasionally stop sets of BCFES trials early in the event of an incompatible finger state (as previously described) or if the decoder consistently predicted a maximum flexion or extension state for at least one trial continuously. This was done in an attempt to avoid the muscles fatiguing too soon into the experiment. In such instances, the decoder was reinitialized and BCFES was re-enabled.

### Patterns

To keep communication bandwidth low, stimulation was delivered by coordinated patterns^28,34^. In a patterned stimulation paradigm, electrodes belonging to a pattern have their pulse parameters governed by a one-dimensional variable, given the range 0-100% in this manuscript. In this way, one command can govern stimulation parameters on any number of electrodes.

In our one-dimensional implementation, we used one pattern to represent stimulation parameters for the electrodes in the FDPi, FDPmrs, and EDC muscles. As implemented on the NNP, patterned stimulation allows one to control pulse width and current amplitude at each command value. Command values were computed by the xPC Target computer from the finger positions predicted by the RKF or the target controller during BCFES and target-controlled FES experiments, respectively. The patterns we implemented did not modulate stimulation amplitude with command value (constant 10mA) but did modulate pulse width between 0 and 255μs. Our pattern delivered maximum stimulation to EDC at 0% command and maximum stimulation to FDPi and FDPmrs at 100% command. In the region around 50% command value, all of EDC, FDPi, and FDPmrs received stimulation at approximately 30% of the maximum pulse width according to the pattern. Generating co-contracting stimulation when not at the limits of range of motion can more consistently create the desired position at that particular command value and combat hysteresis. Supplementary Fig. 4 illustrates an example pattern used during a BCFES experiment.

### Determining Stimulation Command Value

#### Closed-Loop ReFIT Kalman Filter

For all brain-control experiments, we used a ReFIT Kalman filter to predict the intended one-dimensional finger movements^32^. To train the ReFIT Kalman filter, the monkeys first performed at least 325 trials in manipulandum control mode with a 750ms target hold time for Monkey N or 500ms for Monkey W. Then we trained an SBP position/velocity Kalman filter^33^, which Monkey N used in a closed-loop fashion at a 32ms update rate for at least 250 trials with a 500ms hold time. We trained a ReFIT Kalman filter using those closed-loop standard Kalman filter trials and tested it for at least 100 trials. During all brain-control trials, the positions of the fingers as displayed on the screen were integrated from the RKF’s predicted velocities. All of this was performed prior to the experiment’s nerve block, if performed. Following the nerve block, the same ReFIT Kalman filter was used in RKF trials and BCFES trials, where the RKF output was mapped directly to stimulation command value. Manual velocity biases were added as needed to the RKF’s predicted velocities following the nerve block to assist with any changes in noise level resulting from disconnecting and reconnecting the recording hardware.

We implemented a two-finger position/velocity ReFIT Kalman filter in a similar fashion to what we have previously published^31^. The two-finger RKF was never used to control stimulation, as it was limited to 1 degree of freedom control. Training the two-finger RKF followed the same procedure as the one-finger RKF but using two-finger manipulandum control and standard Kalman filter control trials. Following usage of the standard Kalman filter, the ReFIT Kalman filter was trained by rotating the predicted velocity vector towards the single two-dimensional target generated by combining the one-dimensional target for each finger group, similar to the original ReFIT method^53^.

#### Target-Controlled FES

We used target information to control stimulation to better understand the capabilities and challenges when performing continuous functional electrical stimulation. The state machine diagram in Fig. 2a illustrates this controller. At the beginning of each trial, the stimulation command value is set to the target’s center position. Then, a stimulation update was delivered prior to every pulse (32ms between updates) depending on the target’s relative location to the current finger position. If the target required more finger flexion, stimulation command value was increased by approximately 1%. If the target required more finger extension, stimulation command value was decreased by approximately 1%. Updates were sent continuously every 32ms in this fashion until the finger moved to within the middle 75% of the target. Once in the target, stimulation command value was held constant until the finger inadvertently moved out of the middle 75% of the target, the target was successfully acquired, or the trial timeout was reached.

### Performance Metrics

#### Closed-Loop Performance Metrics

We estimated closed-loop performance with success rate and acquisition time, occasionally split into measures of time to target and orbiting time. Success rate was calculated as the total number of targets acquired successfully divided by the total number of targets presented. Acquisition time was computed as the total amount of time from the beginning of the trial to the time the target was successfully acquired and held, less the hold time. Failed trials were given acquisition times equal to the trial timeout less the hold time. Time to target was computed as the time from the beginning of the trial to the first instance the target was touched by the finger. Trials in which the finger never touched its target were given acquisition times and times to target equal to the trial timeout less the hold time. Orbiting time was computed as the time from first touching the target to the time the target was successfully acquired and held, less the hold time. The sum of a trial’s time to target and orbiting time equals its acquisition time.

In all metrics, the following conditions were excluded from analysis to avoid biasing the results. Successful trials immediately following a failure were excluded to avoid situations in which the monkey would wait at the central position to acquire the central target once it returned following a failed outer target. Trials immediately following a change in control method were also excluded, as the state of the controller could not be determined prior to usage. For example, when switching from RKF control to manipulandum control, the displayed finger position could jump from outside of the target straight to the subsequent target, resulting in an unrealistically small acquisition time.

For the orbiting time metric, trials in which the finger never touched its target were excluded, as orbiting time could not be computed.

#### Open-Loop Performance Metrics

We estimated open-loop prediction performance using the intra-class correlation coefficient. As opposed to Pearson’s correlation, ICC is a better measure of agreement between two conditions. ICC is computed via the following equation (1):

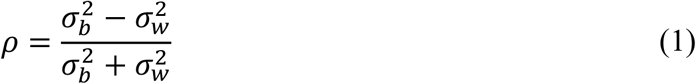

where 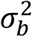 is the variance between conditions and 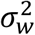 is the variance within conditions. Confidence intervals were computed via the following formulas and equation (2)^54^:

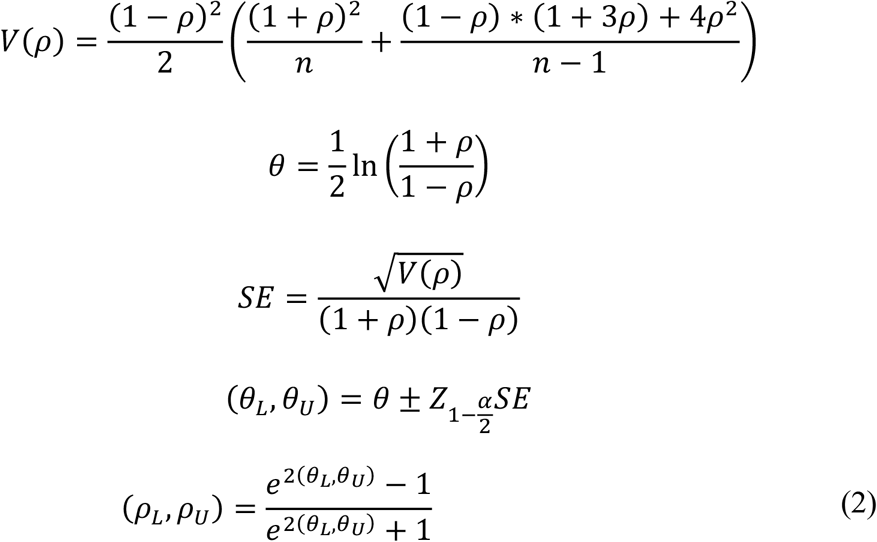

where 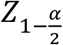 is the *z*-score at the confidence level 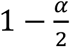 Significant differences between ICCs were determined via the Fisher’s transformation of *θ* for each ICC via equation (3):

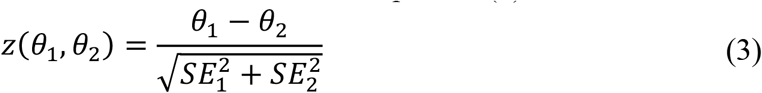

The *p*-value for *z* was calculated from a normal distribution.

## ACKNOWLEDGMENTS

We thank Gail Rising, Amber Yanovich, Lisa Burlingame, Patrick Lester, Veronica Dunivant, Laura Durham, Taryn Hetrick, Helen Noack, Deanna Renner, Michael Bradley, Goldia Chan, Kelsey Cornelius, Courtney Hunter, Lauren Krueger, Russell Nichols, Brooke Pallas, Catherine Si, Anna Skorupski, Jessica Xu, and Jibing Yang for expert surgical assistance and veterinary care. We appreciate Chris Andrew’s expert statistical assistance. This work was supported by NSF grant 1926576, Craig H. Neilsen Foundation project 315108, A. Alfred Taubman Medical Research Institute, NIH grants R01GM111293 and R01EB024522, and MCubed project 1482. S.R.N-T¶. was supported by NIH grant F31HD098804. M.J.M. was supported by NSF grant 1926576. E.K. was supported by ___. J.M.L. was supported by NIH grant R01EB024522. K.L.K. was supported by NSF grant 1926576 and NIH grant R01EB024522. T.A.K. was supported by NIH grant R01NS105132. M.S.W. was supported by NIH grant T32NS007222. P.G.P. was supported by NSF grant 1926576, A. Alfred Taubman Medical Research Institute, and NIH grant R01GM111293. C.A.C. was supported by NSF grant 1926576, Craig H. Neilsen Foundation project 315108, NIH grant R01GM111293, and MCubed project 1482.

## AUTHOR CONTRIBUTIONS

C.A.C., P.G.P., and M.S.W. advised the work and performed the cortical surgeries. T.A.K. and N.G.K. performed the upper extremity surgeries. P.G.P., S.C., and E.K. designed and performed the nerve blocking procedure. K.L.K. and J.M.L. advised FES and NNP work. M.J.M. and S.R.N-T. conducted experiments, collected data, and wrote and executed code on the data. S.R.N-T. wrote the manuscript and led the experiments. All authors reviewed, edited, and approved the manuscript.

## DATA AVAILABILITY

A subset of the data used in this manuscript will be available online following publication. The full set of data will be available from the corresponding authors upon reasonable request.

## CODE AVAILABILITY

Code will be available from the corresponding authors upon reasonable request following publication.

