## Supplementary Information for "Brain-Controlled Electrical Stimulation Restores Continuous Finger Function"

[1]Department of Biomedical Engineering, University of Michigan, Ann Arbor, MI 48109, United States of America
[2]Department of Biomedical Engineering, Case Western Reserve University, Cleveland, OH 44106, United States of America
[3]Department of Orthopaedics, MetroHealth Medical Center, Cleveland, OH 44106, United States of America
[4]Louis Stokes Cleveland Department, Veterans Affairs Medical Center, Cleveland, OH 44106, United States of America
[5]Department of Anesthesia, University of Michigan Medical School, Ann Arbor, MI 48109, United States of America
[6]Section of Plastic Surgery, Department of Surgery, University of Michigan, Ann Arbor, MI 48109, United States of America
[7]Department of Neurosurgery, University of Michigan Medical School, Ann Arbor, MI 48109, United States of America
[8]Department of Electrical Engineering and Computer Science, University of Michigan, Ann Arbor, MI 48109, United States of America
[9]Robotics Institute, University of Michigan, Ann Arbor, MI 48109, United States of America
[10]Neuroscience Graduate Program, University of Michigan, Ann Arbor, MI 48109, United States of America
[11]Department of Neurology, University of Michigan Medical School, Ann Arbor, MI 48109, United States of America
[†]These authors contributed equally to this work.

### SUPPLEMENTARY METHODS

#### Controller for Target-Controlled Functional Electrical Stimulation

To perform target-controlled functional electrical stimulation, we delivered intramuscular stimulation according to the state machine illustrated in Supplementary Fig. 1.

#### Cortical Implants

To perform brain-controlled functional electrical stimulation for the restoration of continuous hand function, we implanted Monkey N with three Utah microelectrode arrays in each of the hand areas of left and right hemisphere primary motor and sensory cortices and Monkey W with two Utah microelectrode arrays in the hand area of left hemisphere primary motor and sensory cortices. Supplementary Fig. 2 presents photographs of the implants.

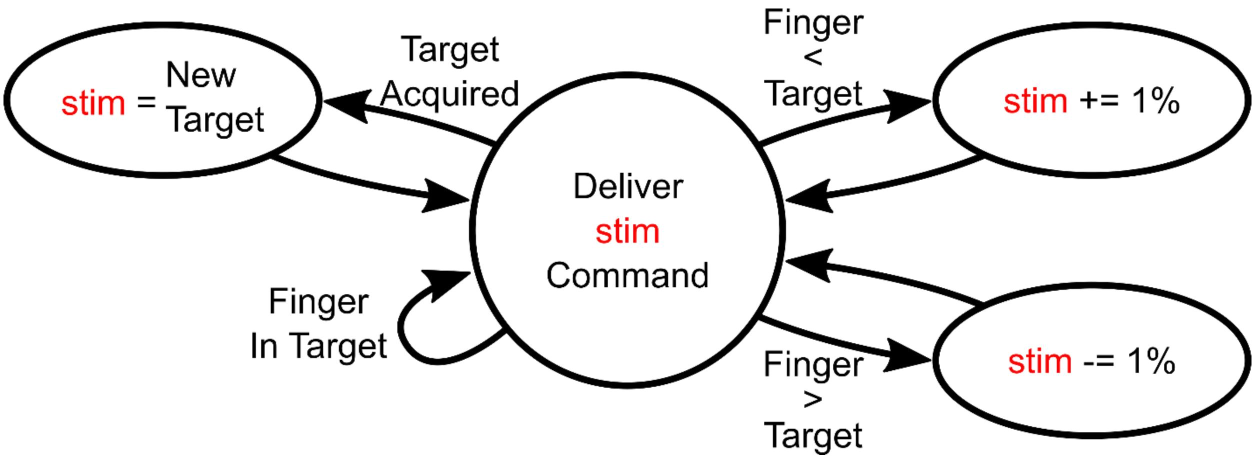


**Supplementary Fig. 1 State machine diagram illustrating the control system used to deliver target controlled stimulation.**

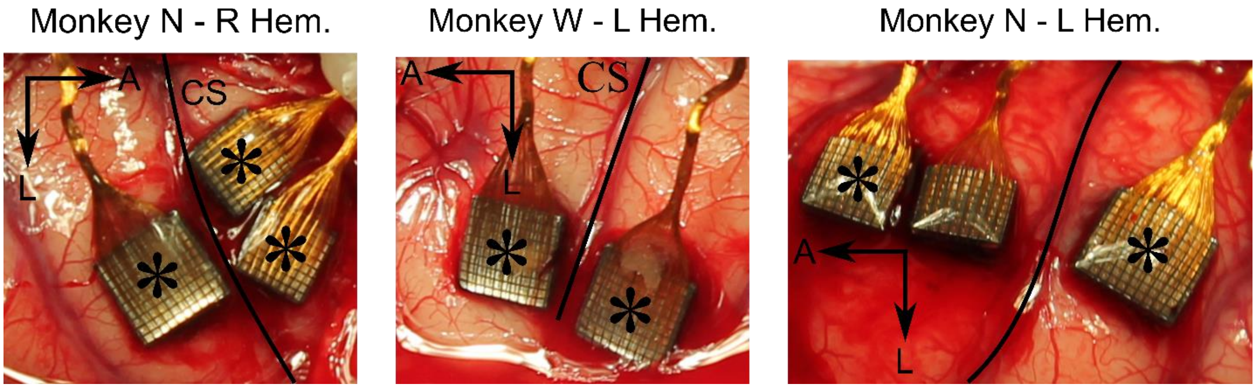


**Supplementary Fig. 2 Photograph of Monkey N's and W's Utah microelectrode array implant locations.** Larger units are 10x10 arrays with 96 active electrodes. Smaller units are 8x8 arrays with 64 active electrodes each.

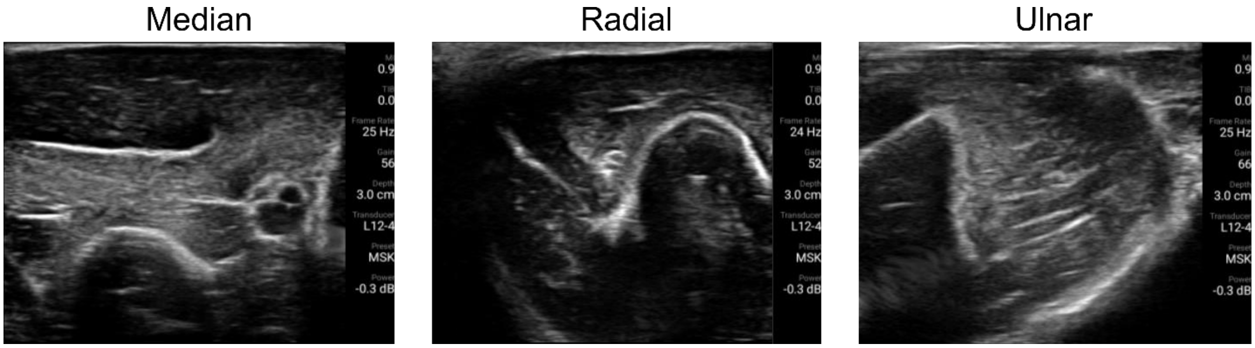


**Supplementary Fig. 3 Example ultrasound images taken during a nerve block procedure.** All images were captured along a transverse plane just proximal to the elbow with Monkey N.

#### Ultrasound Examples

To guarantee that the monkey's capable movements were purely a result of the BCFES system and not the monkey's native efforts, we delivered nerve blocks to the median, radial, and ulnar nerves of his left arm to temporarily block signal transduction to the arm. We used ultrasound to deliver a 2% lidocaine with 1:100,000 epinephrine to the tissue surrounding the nerves to block activity. Supplementary Fig. 3 displays some example images captured from one experiment.

#### Functional Electrical Stimulation Pattern

To deliver electrical stimulation in a low-bandwidth communication space, we used stimulation patterns to govern parameters for all electrodes innervating the hand. The controller commanded a value on the horizontal axis in Supplementary Fig. 4, and the Networked Neuroprosthesis commanded the corresponding pulse widths for each of the active electrodes.

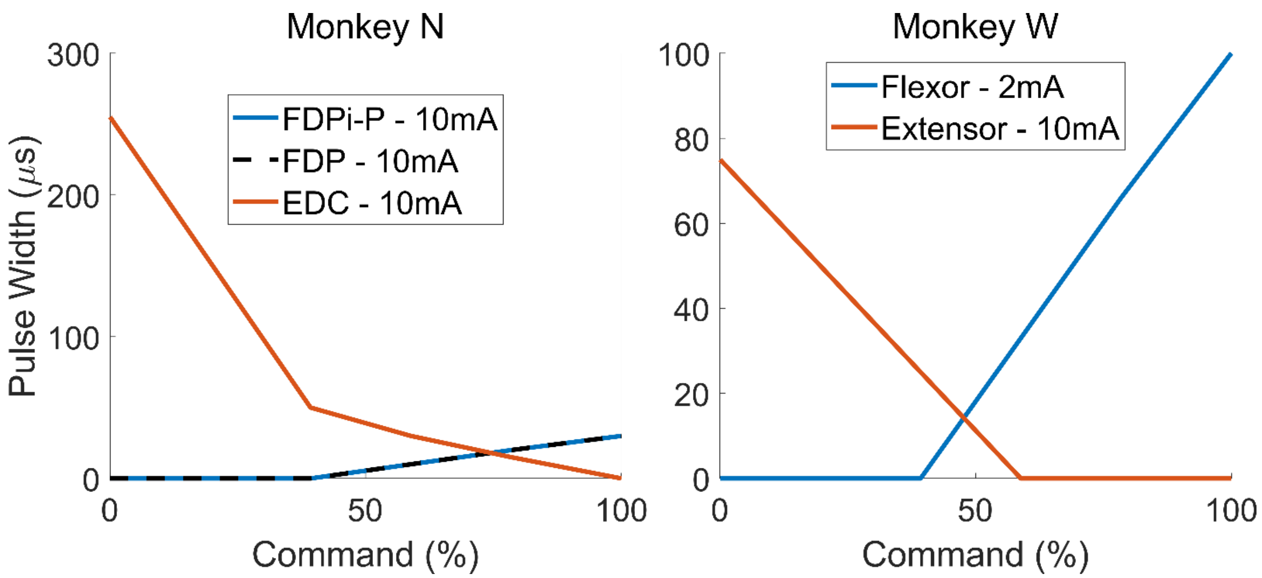


**Supplementary Fig. 4 An example stimulation pattern used during a brain-controlled functional electrical stimulation experiment with Monkey N or a target-controlled functional electrical stimulation experiment with Monkey W.**

### SUPPLEMENTARY ANALYSIS

#### Brain-Controlled FES Performance May Be Improved with Lower Latency

Muscle fatigue and hysteresis may increase the difficulty of acquiring targets, particularly with simple controllers like the one investigated in Supplementary Fig. 1. In the case of any feedback controller, latency in the feedback loop can also increase difficulty by introducing ringing (or orbiting as we refer to it in the context of target acquisition) without proper dampening. We next sought to investigate the impact of controller latency on Monkey N's capability of controlling the virtual hand using able-bodied manipulandum control and the BCFES system.

We estimated the relative latencies of our various control methods for the 1D task. To do so, we offset the measured or predicted behavior and SBP from -200ms (behavior precedes cortical activity by 200ms) to +400ms (cortical activity precedes behavior by 400ms) in 1ms steps and used ridge regression ($\lambda = 0.0001$) to predict the movement velocities at each offset. The optimal latency for each control mode was calculated as the maximum intra-class correlation across all offsets, as fit by a parabola around the true maximum intra-class correlation. We compared latencies of three implants in three cortical hemispheres in two monkeys, historical data from Monkeys N (left hemisphere) and W in our past publication (Nason et al., 2020) and Monkey N data from this publication, which used right hemisphere.

In all cases, we found reasonable latencies between cortical SBP and behavioral outputs on the order of 22.75 to 41.5ms ($25^{th}$ and $75^{th}$ percentiles, respectively, $n = 25$) in manipulandum control mode. Note these estimates do not take into consideration the processing and communication latencies of the BMI hardware, so are not necessarily representative of true spinal cord transduction speeds but rather can be compared relatively. Supplementary Fig. 5a presents histograms illustrating the optimal latencies of all datasets analyzed for each implant. Data from Monkeys N and W from our past work show similar latencies between 25.75 and 70.5ms ($25^{th}$ and $75^{th}$

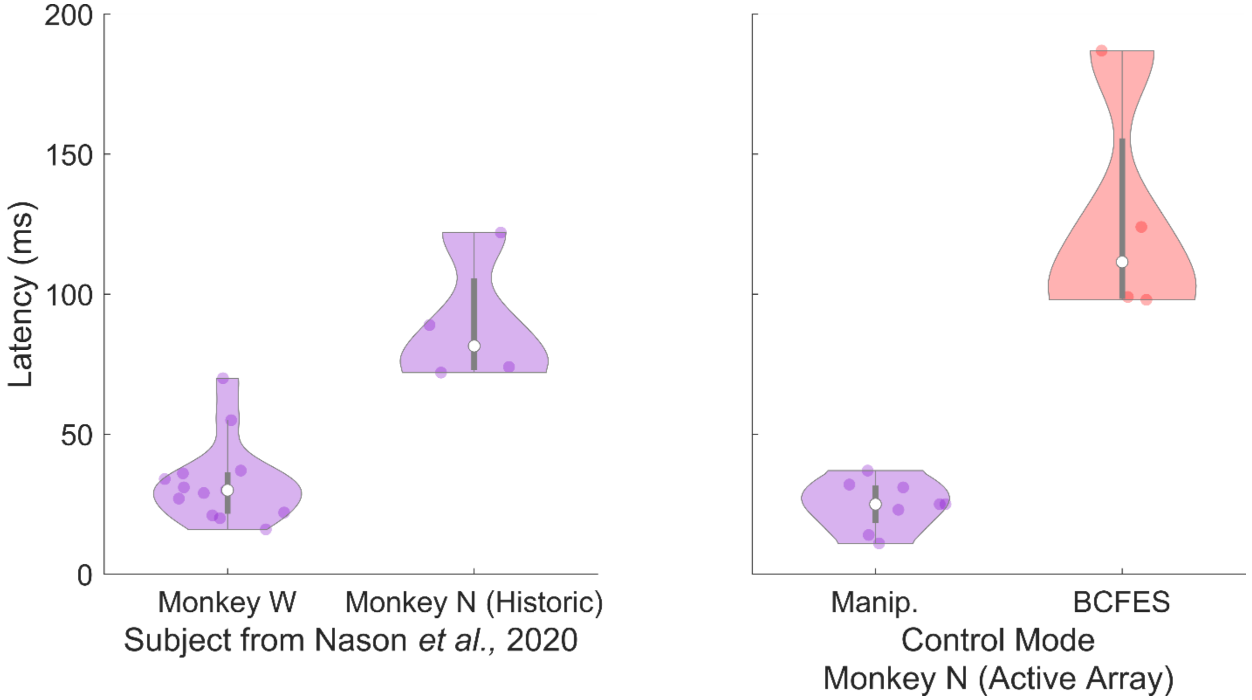


**Supplementary Fig. 5 Latency of BCFES compared to manipulandum control.** (**a**) Violin plots illustrating optimal latencies for various implants in various control settings. Each colored dot represents one day of recording. The white dot represents the median and the gray bar represents the 25$^{th}$ and 75$^{th}$ percentiles. (*Left*) Optimal manipulandum control latencies for Monkeys W and N from Nason *et al.*, 2020. (*Right*) Optimal latencies for Monkey N using the manipulandum and the BCFES system.

percentiles, respectively, $n = 17$), though Monkey N's RKF latencies were higher likely due to the more anterior placement of the recording array. Monkey N's latencies from this work were very consistently between 18.5-31.5ms (25$^{th}$ and 75$^{th}$ percentiles, respectively, $n = 8$). At their optimal latencies, we found that the peak velocity intra-class correlations were reasonably high for single time-step ridge regressions at 0.15 to 0.38 (25$^{th}$ and 75$^{th}$ percentiles, respectively, $n = 25$).

When Monkey N used the BCFES system, we found that the optimal latency between cortical activity and the resulting FES-controlled behavior was substantially higher at 98.5 to 155.5ms (25$^{th}$ and 75$^{th}$ percentiles, respectively, $n = 4$), despite the peak velocity intra-class correlations staying reasonable at 0.15 to 0.23 (25$^{th}$ and 75$^{th}$ percentiles, respectively). The substantial increase in latency compared to manipulandum control is likely due to the wireless transmission of stimulation commands, real-time computation of stimulation updates in the embedded hardware, and generation of a contraction response in the target musculature.
